# Overlapping flower and pollen production in two imperiled pitcher plants raises conservation challenges

**DOI:** 10.64898/2026.09.02.748929

**Authors:** Rebecca E. Hale, Caroline Kennedy, Wayne Morgan, Todd Brasseur, Elizabeth Companion, William Gay, Kristen Hillegass, Alyssa Lynch, Michelle Paredes, Gabi Parker, Lila Uzzell, Mars Zappia, Jennifer Rhode Ward

## Abstract

Balancing sexual reproduction with vegetative growth poses challenges for many plants. Additionally, reproductive effort, such as production of pollen or flowers, is not always correlated with numbers of viable and germinable seeds. Energetic tradeoffs like these can be particularly fraught for imperiled species and might be especially complicated in landscapes with a possibility of interspecific hybridization. In this multi-year study, we track the reproductive effort, flowering phenology, and reproductive output of two interfertile *Sarraceni*a (pitcher plants). We show that reproductive effort varies among years and sites, and that production of pollen and flowers does not always result in higher seed output. We demonstrate that phenological overlap in the timing of flower production and pollen viability could enable hybridization, and that the more imperiled taxon produces fewer seeds per flower. Results have implications for conservation of these pitcher plants, management of hybridization in these and other systems, and general principles of reproductive allocation.

## Introduction

Resource allocation theory describes energetic tradeoffs that balance organisms’ maintenance, growth, defense, and reproduction (1, 2). Reproductive effort is the proportion of an organism’s bulk energy committed to its reproductive organs and gametes (3). For angiosperms, this is the energy diverted to producing flowers and gametes (4, 5), often quantified as pollen and ovule numbers and ratios (6). An organism’s reproductive effort can be correlated with its reproductive output (3). For angiosperms, this is commonly estimated as seed quantity and/or quality (2), which can be limited by pollen quality, pollen quantity, and pollinator activity (7, 8, 9). Both reproductive effort and output can also be allometrically constrained (10). Fully characterizing plants’ reproductive effort and output, and related costs, is challenging and often difficult to predict (11) but might be particularly important to the conservation of imperiled taxa (12).

The ways in which angiosperm reproductive effort and output manifest are subject to tradeoffs. For example, in a study of three *Vallisneria* species, which rely heavily on asexual reproduction, species with more flowers per ramet also produced more seeds per flower (13); this could reflect taxonomic differences in allocation to vegetative versus reproductive tissue. A review of 49 taxa found that larger flowers made more and larger ovules in actinomorphic species (14), and a metaanalysis of 69 monocot families showed that larger flowers tended to produce more seeds (15). However, in Orchidaceae, relationships among reproductive traits are more complex. Single-flowered species were more likely to have pollinaria removed by visitors and to produce larger seeds, but multi-flowered species were more likely to fruit (16). This study suggested that ways in which reproductive effort translates to reproductive output can depend on the behavior of animal visitors and subsequent pollination success. These tradeoffs can be influenced by surrounding plant communities (e.g., 17) or abiotic environmental factors (e.g., 18).

Divergent allocation to reproductive structures among closely related species may serve to reduce hybridization through pre-zygotic reproductive isolation. In numerous animal pollinated species, pre-zygotic barriers, including ecogeographic isolation, phenology, pollinator behavior, and pollen precedence, are more important drivers of reproductive isolation than post-zygotic mechanisms (19, 20, 21). Pre-zygotic reproductive isolation can also be facilitated by low total investment into sexual reproduction by species with significant use of vegetative strategies (e.g., *Vallisneria* spp., *Fragaria* spp., and *Sarracenia* spp.). Such pre-zygotic barriers reduce energy allocation to hybrid seeds, which might have lower viability or fertility (e.g., 20, 22).

The southern Appalachian’s mountain bogs are habitat for numerous imperiled species of animals and plants (23, 24), including two rhizomatous, rosette-forming carnivorous taxa in the Sarraceniaceae (Pitcher Plant) family. These pitcher plants occupy broadly overlapping ranges and naturally co-occur at six distinct sites in North and South Carolina. In these habitats, where nutrient availability is low, the plants’ tubular leaves lure, capture, and digest insects as supplemental nitrogen sources (25). Both taxa are federally listed in the U.S., mostly due to habitat loss and poaching (26, 27).

*Sarraceniaceae* flowers are self-compatible with no obvious dichogamy (28), and they typically outcross (29). Selfing results in lower quality and quantity of offspring (30), and the style’s shape discourages self-pollination within a flower (31). Each species produces downward-oriented, complete, actinomorphic flowers that hang pendulously from the stem; flowers typically have five sepals, five petals, over 80 stamens, and five carpels (31, 32). Nectar is produced in the ovary walls and extraflorally (33). Specific pollinators are poorly described but could include bumblebees (*Bombus*) and sarcophagid flies (25). In a study of one *Sarracenia* species at two New England bogs, availability of nitrogen (from prey capture) and carbon (from photosynthesis) seemed to influence seed production more than pollen limitation (25).

While the two taxa examined in this study are separated by nearly 4 my (34), they can produce fertile hybrids when grown in sympatry (35, 36). Hybridization like this, involving one or more imperiled congeners, can pose conservation challenges (37). The occurrence of phenotypic hybrids in many natural sites could indicate that both pre- and post-zygotic barriers to hybridization are inadequate to prevent interspecific fertilization. The purpose of this study was to quantify the reproductive effort and output of two *Sarracenia* from populations across western North Carolina, while evaluating divergent phenology as a mechanism of pre-zygotic reproductive isolation. We did this across multiple years and included sites where *Sarracenia purpurea* var. *montana* is the only member of its genus along with sites where it co-occurs with *Sarracenia rubra* ssp. *jonesii*. Understanding reproductive differences between these taxa could inform conservation efforts and be used to manage hybridization potential.

## Methods

### Study Species

The Mountain Purple Pitcher Plant (*Sarracenia purpurea* var*. montana* D.E. Schnell & Determann) is restricted to western North Carolina, northern South Carolina, and northern Georgia. *S. purpurea* var*. montana* produces short, decumbent pitchers with flowers that grow as high, or higher, than its modified leaves. Leaves’ distal hood tips tend to curve inwards rather than flare outwards like its close relatives (38, 39). The species has low, broad pitchers that grow laterally and range in length from 6.5 - 18 cm. Their scapes are erect, with a single terminal flower. Scape height ranges from 26.5 - 64.5 cm (40; T. Brasseur, unpublished data). Habitat loss and poaching have rendered the Mountain Purple Pitcher Plant a critically imperiled taxon (T1; NatureServe 2026), and it is under review for federal listing (ECOSphere Environmental Conservation Online System 2024).

The taxonomic ranking of Mountain Sweet Pitcher Plant, *Sarracenia rubra* ssp. *jonesii* Wherry (Wherry), as a species or subspecies is unresolved; this paper utilizes the taxonomy of the U.S. Fish and Wildlife Service and Integrated Taxonomic Information System. This taxon is found only in North and South Carolina. It has erect, slender pitchers that range in height from 21 - 73 cm (39, 41). The taxon produces fragrant flowers on scapes 32 - 70 cm (42) that often grow higher than its modified leaves (43). *S. rubra* ssp. *jonesii* is globally imperiled (G2; NatureServe 2026) and has been listed as federally endangered since 1988 (41).

### Study Areas

Sites are referred to by abbreviations (Table 1) to obscure geographic locations of these imperiled taxa and reduce the incidence of poaching. In 2015 we determined total seed production of one or both species at six sites (CM, DB, HC, MB, RL, SF). From 2019 - 2023, both species were monitored across four sites at which they naturally co-occurred (CM, MB, RL, SV) to assess reproductive effort (flower production, pollen production, flower phenology) and output (seed production). Due to the labor-intensive nature of the data collection, we were only able to sample two sites per year.

**Table 1.**
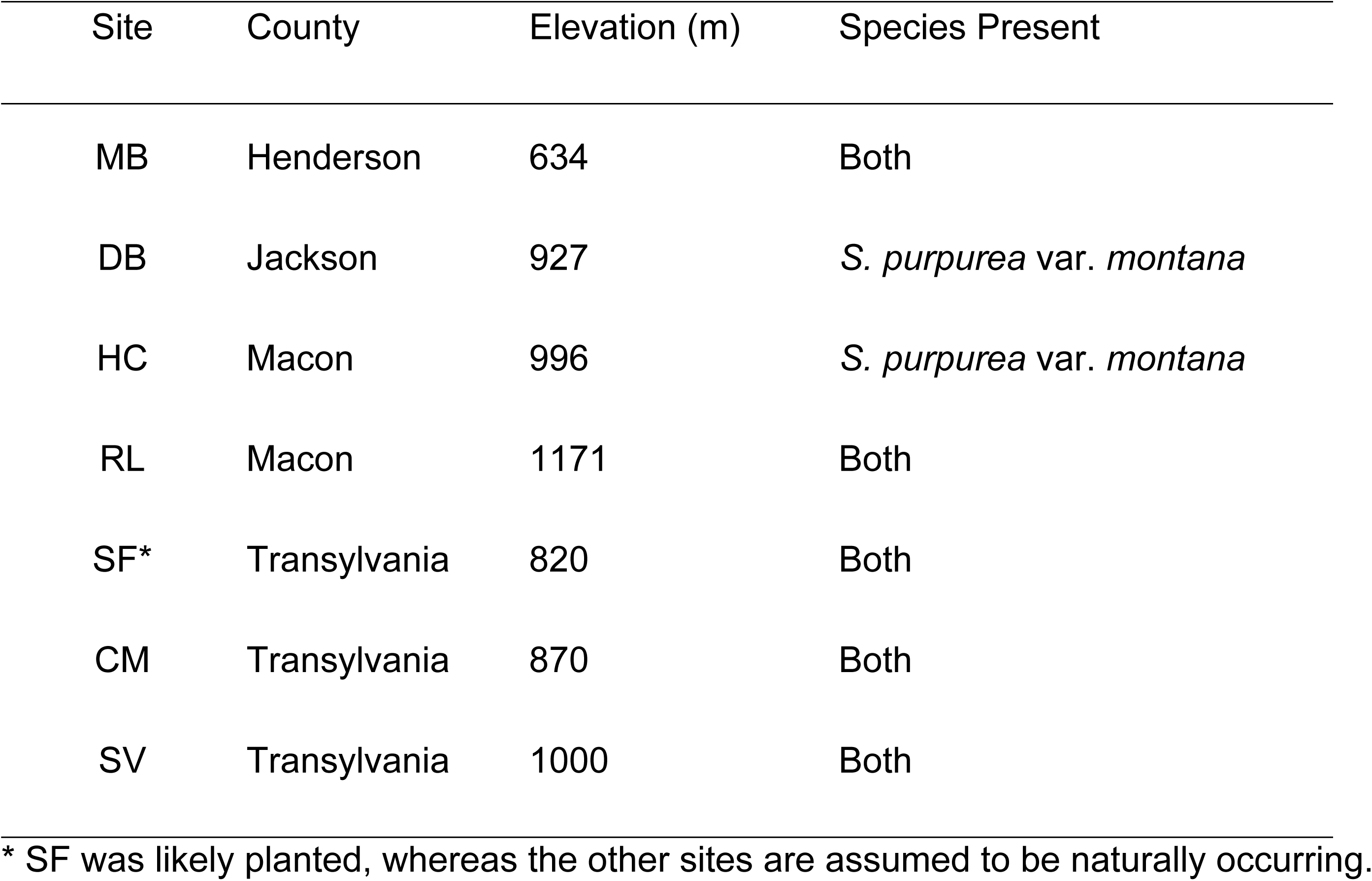
Site identity, location by county, and elevation for the seven sites included in the study.

Sites are identified by two-letter codes to obscure their geographic locations. All sampling was carried out under permits from the North Carolina Plant Conservation Program (permit #s 456, 600, 638, 708, 774, 831, 889, 945) and the North Carolina Forest Service (FM23015). In addition, we obtained written permission to work on private property.

Because pitcher plants grow in networks of rosettes, identifying individual ramets can be challenging. The U.S. Fish and Wildlife Service classifies ramets as distinct clumps when separated by at least 0.5 m (44), and we adopted this convention. Twenty clumps per taxon and site were sampled and, in sites with fewer than 20 clumps of one or both taxa, all were sampled. At one site and year (SV, 2023), every (64) *S. purpurea* var*. montana* clump was sampled along with every (9) *S. rubra* ssp*. jonesii*.

### Flower Production

#### Data collection

All flowers on our focal clumps, including buds and those that were fully open, were counted in May of each sampling year. Although clumps continued to produce flowers through summer, we assumed, based on previous observations, that this early- season flower production was representative of production over the season. Sites were visited weekly until all marked flowers had senesced (stage 6).

#### Data analysis

A large proportion of clumps of both species did not produce flowers. Therefore, we first converted flower counts to a binary variable (flowers/no flowers). Differences in the odds of producing flowers among sites and between species were evaluated using logistic regression with the glm function, specifying binomial error, in R (45).

Determination of significance in this an all analyses applied an *α* = 0.05. Effects of individual predictors were obtained using likelihood ratio tests implemented with the drop1 function, starting with a model including the predictors of site, species, and their interaction, then removing the interaction to evaluate main effects. Since interannual variation in flower count was not the focus of our study, we pooled data for each site across years. After evaluating the odds of producing flowers, we then subset only those clumps that produced at least one flower and examined whether flower count differed between species or among sites. Flower count distributions were non-normal in each species; therefore, we performed a Scheirer-Ray-Hare test on the number of flowers produced.

### Flower Stage and Pollen Viability

#### Data collection

Beginning in 2019, we quantified flowering phenology and reproductive output for both species. We selected up to three flowers per clump and scored each flower’s stage (Table 2). Floral stage categories were defined because they were easy to quantify in the field and may capture important transitions in the life stages of these flowers. We collected one stamen per flower weekly from mid-May until focal flowers had senesced. While reproductive effort is best measured by assessing all reproductive structures (including pollen, ovules, sterile floral whorls, and nectar; 46), we measured only viable pollen production to avoid destructive sampling of imperiled plants.

**Table 2.**
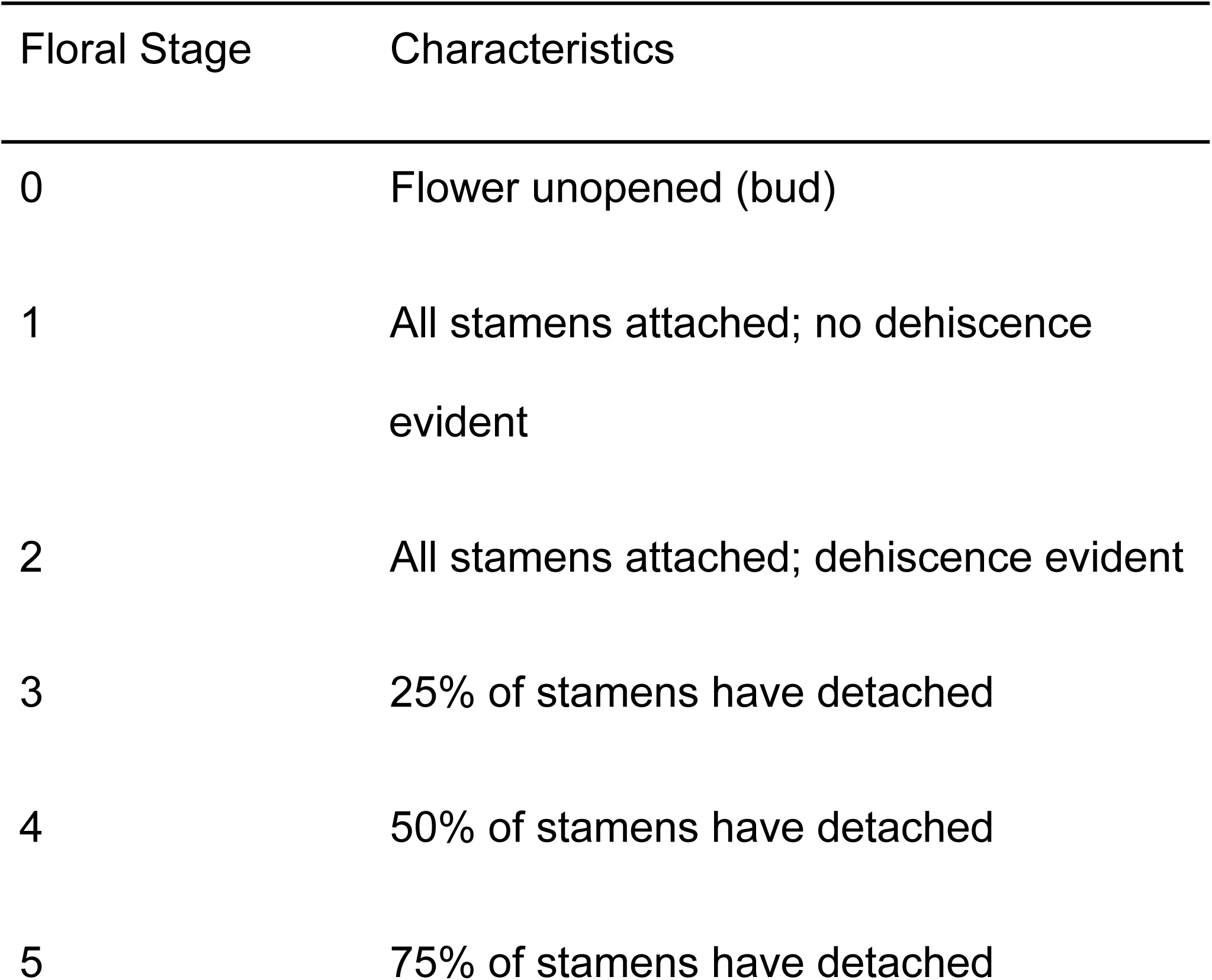
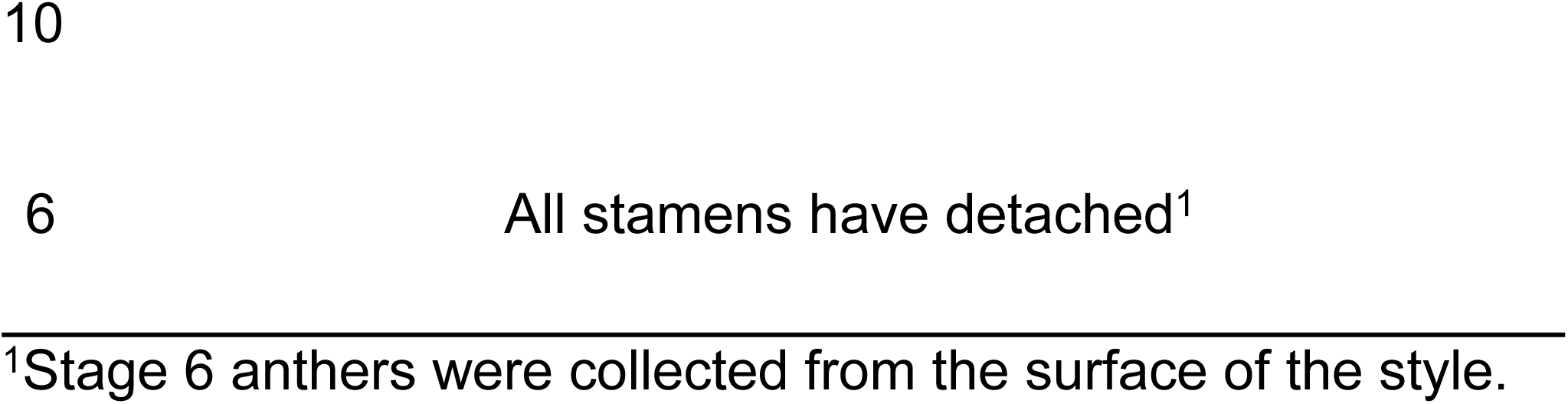
Criteria for distinguishing floral development stage.

We preserved and stained anthers using methods from Kearns and Inouye (47). To quantify the amount of pollen per anther, we added 10% (10 μL) of the sample onto a hemocytometer with Neubauer rulings. Viable and inviable pollen were counted under an Olympus CH30 compound microscope at 400X total magnification, using a hemacytometer with a 0.9 µl * 9 grid.

We calculated the total anther pollen, P_T_, as

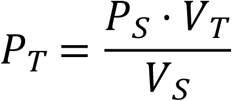

where *P_S_*was the pollen counted in our sample, *V_T_* was the total volume of the full suspension (100 µl), and *V_S_* was the sample volume (0.9 µl).

#### Data analysis

Across four sites, from 2019 to 2023, a total of 729 flowers were sampled for flower stage and 574 for pollen viability across all sites. Since flowers often developed through multiple stages during the one week sampling interval, not all flowers were sampled in each stage.

Although up to three flowers per clump were sampled, all flowers were treated as independent subjects. We did not consider clump identity in our analyses. The proportion of flowers in each stage over time was compared between species using cumulative link mixed models assuming proportional odds (no interaction between fixed effects), implemented with the ordinal package in R (48). The variable “flower identity” was treated as a random effect to account for repeated sampling of flowers over time. Fixed effects of date and species were evaluated with likelihood ratio tests using the *drop1* function. Because the phenology of bud production varied across sites and years, each site-year combination was evaluated separately.

To evaluate how viable pollen quantities changed across flower stages, we performed Spearman rank correlations separately for each species, treating stage as an ordinal independent variable. Sites and years were pooled, as previous studies of these populations found no differences among sites or years in the relationship between the amount of viable pollen and flower stage (49, 50). Repeated samplings of the same flowers were treated as independent, which could artificially inflate any correlation.

To compare pollen production between species, we evaluated pollen counts in stage 2 flowers, only. Shapiro-Wilk normality tests on pollen counts (both logged and not logged) for each site-species combination and found non-normal distributions for one site. Therefore, we performed a Scheirer-Ray-Hare test on the number of pollen grains produced. For *post hoc* comparisons, we used Mann-Whitney U tests and decided *a priori* to evaluate differences in pollen production between species at each of the four sites (years pooled; four tests total).

### Seed Production

#### Data collection

A total of 232 flowers were sampled for seed production across 7 sites between 2015 and 2023 (S1 Table). SF was excluded from the study after 2015 because it contains individuals of unknown provenance and *Sarracenia* species not native to the region.

At each site, mature ovaries were collected between late September and mid-October by placing a paper bag over the flower, pinching the opening closed around the scape, and cutting the base of the scape. Samples were stored in open bags at room temperature until dry, then the outer layers of petals and sepals were removed. Seeds and unfertilized ovules were removed from each ovary’s five locules, and seeds were counted under an Olympus SZ51 Stereo Zoom microscope at 4-10X total magnification. After counting, seeds were accessioned into the North Carolina Botanical Garden collection or returned to their site of origin, in accordance with the conditions of the relevant collection permits. A subset of RL seeds was retained for germination studies.

### Data analysis

A large proportion of sampled flowers did not produce seeds, so we first converted seed counts to a binary variable (seeds/no seeds). Differences in the odds of producing seeds among sites and between species were evaluated with logistic regression using the *glm* function in R, specifying binomial error. Site and species effects were evaluated for significance using likelihood ratio tests implemented with the *drop1* function, starting with a model including the main effects of site, species, and their interaction, and then removing the interaction to evaluate main effects. Since interannual variation was not the focus of our study, we pooled seed production for each site across years.

Seed data were then subset to include only flowers that produced seeds and only the four sites that were the focus of our 2019-2023 sampling: CM, MB, RL, and SV. Shapiro-Wilk normality tests on seed count (both logged and not logged) for each site-species combination revealed non-normal distributions for some sites in each year. Therefore, we performed a Scheirer-Ray-Hare test on the number of seeds produced using the rcompanion package of R (51). For *post hoc* comparisons, we used Mann-Whitney U tests and decided *a priori* to evaluate differences in seed production between species at each of the four sites (years pooled; four tests total).

### Seed Germination

#### Data collection

Five seeds from each RL plant sampled in 2021 (n = 12 *S. purpurea* var*. montana*; n = 11 *S. rubra* ssp. *jonesii* individuals) were weighed individually with a Mettler Toledo XP2U Ultra-Micro Balance scale (± 0.001 mg). Different quantities of the two species were sampled due to permit restrictions and seed availability limitations.

A different group of 10 seeds from 23 RL flowers were prepared for lab germination. Although mass was not measured for the seeds that were subsequently allowed to germinate, it was measured for seeds from the same fruit. These seeds were surface-sterilized using a 10% bleach solution, triple-rinsed in sterilized deionized water, and arranged in a 90 mm Petri dish between two autoclaved filter paper disks saturated with 6 mL of sterile deionized water. Seeds were then stratified at 4°C for five weeks (52). Following the five-week period, Petri dishes were placed on a lab benchtop and exposed to sunlight filtered through a window and ambient conditions (temperature 20.5°C–21.9°C, humidity 33%–47%). Every two days for up to 46 days, seeds were examined with an Olympus SZ51 Stereo Zoom microscope (7–30X total magnification) for radicle emergence and were supplemented with an additional 2 mL of sterilized deionized water.

#### Data analysis

Seed mass was compared among species using a linear mixed effect model, in which seeds were nested within plant clumps, using the *lmer* function of the lme4 package in R (53). Germination success was evaluated with logistic regression using the *glm* function of R, specifying binomial error. The effect of species was determined using the *drop1* function. We evaluated whether differences in germination rate could be explained by differences in seed mass using models in which average seed mass and species were included as independent variables.

## Results

### Flower production

The percent of clumps with flowers ranged from 50% to 95% across sites and species (Fig 1), and the number of flowers per clump was highly variable (mean ± SD: 5.50 ± 10.86; Fig 2). Probability of producing flowers did not differ between species or among sites (likelihood ratio test; species: X^2^ = 0.16, p = 0.69; site: X^2^ = 1.76, p = 0.62). However, among those that produced flowers, *S. rubra* ssp. *jonesii* clumps produced significantly more flowers than *S. purpurea* var*. montana* clumps (Scheirer Ray Hare test; species x site interaction: H = 2.09, df = 3, p = 0.55; species: H = 29.16, df = 1, p < 0.001; site: H = 12.96, df = 3, p = 0.0047).

**Fig 1.**
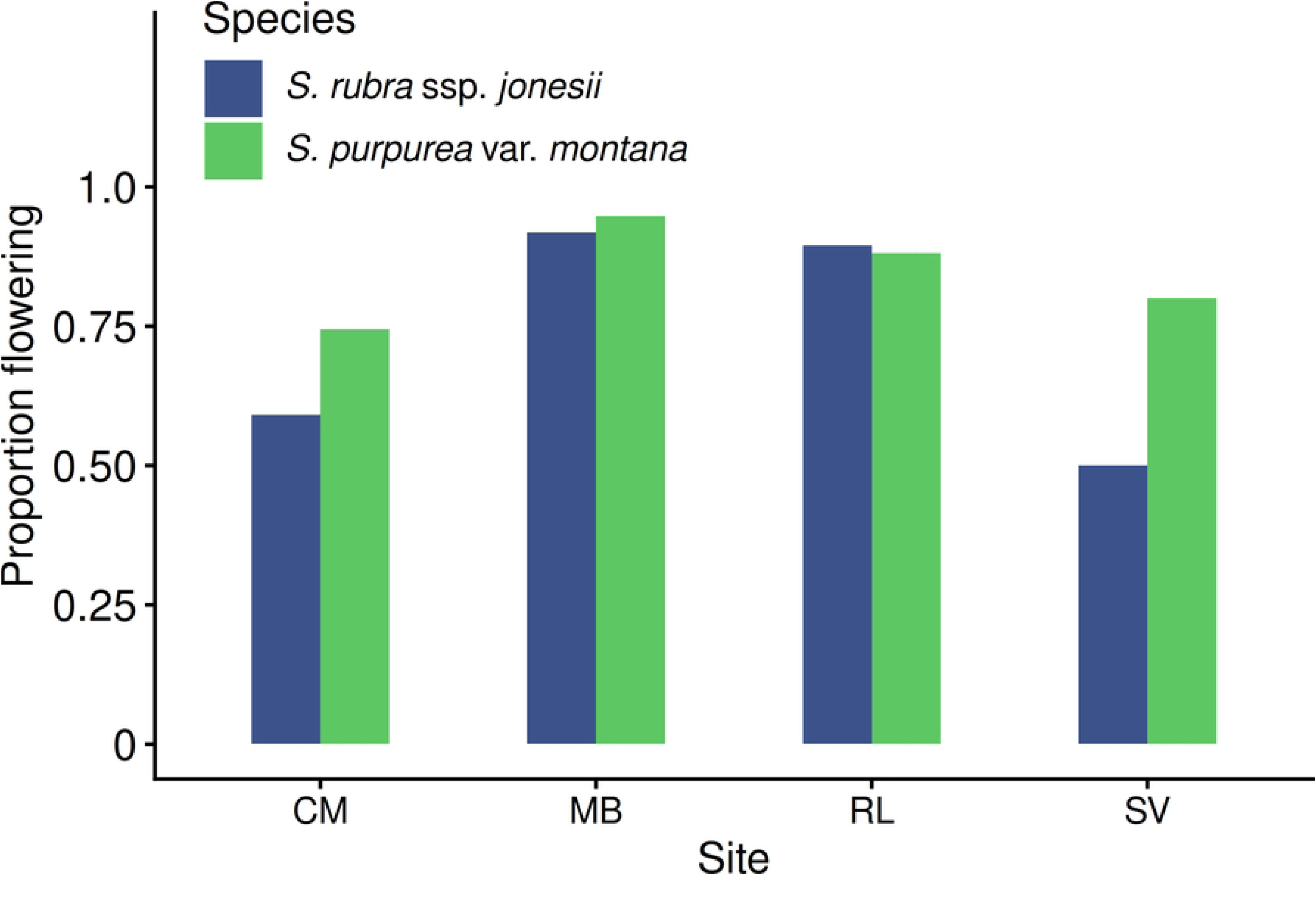
The proportion of *S. rubra* ssp. *jonesii* and *S. purpurea* var. *montana* plant clumps that produced flowers at each of four sites. Plants were sampled across multiple years but are pooled here and in the analyses.

**Fig 2.**
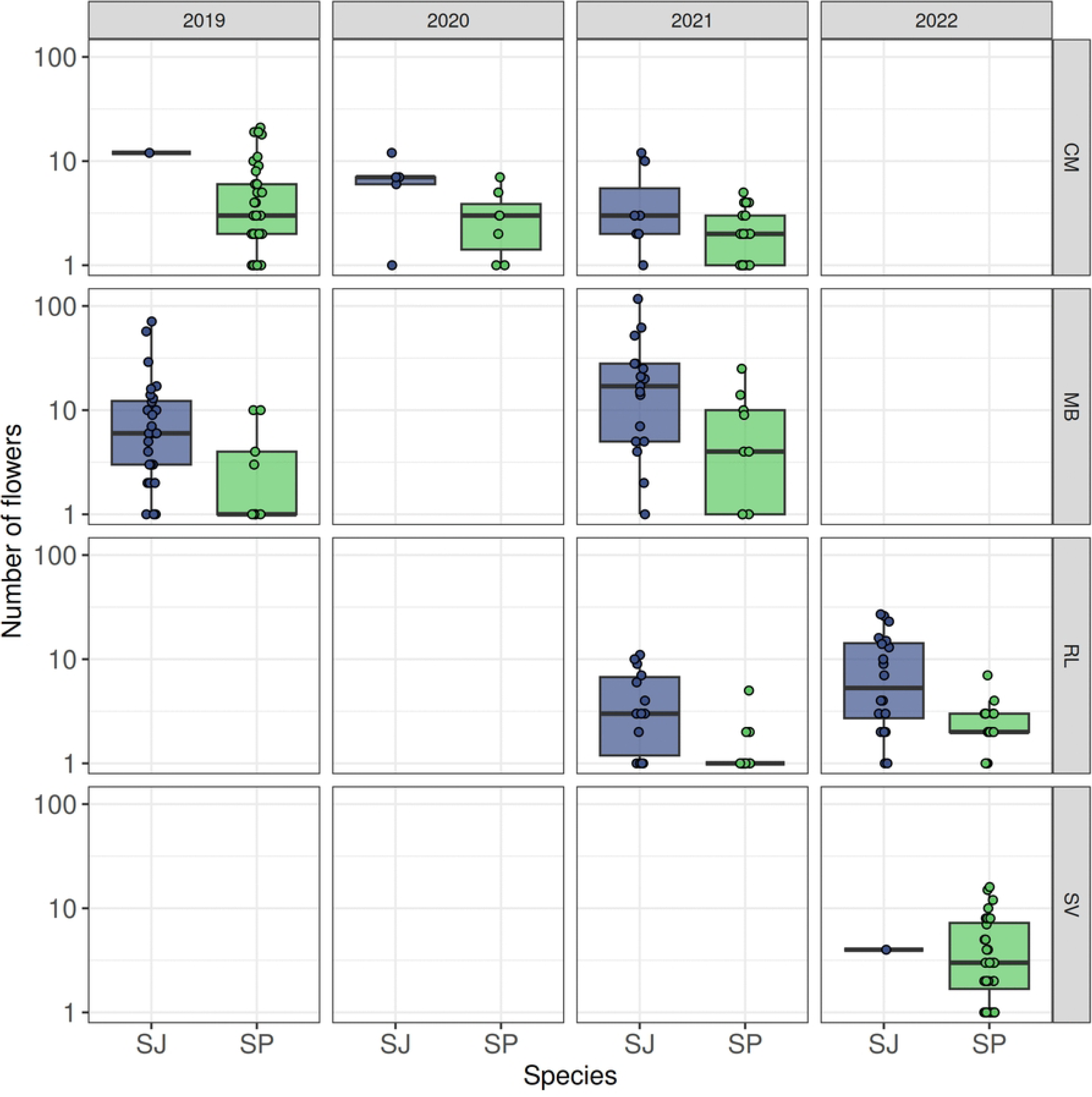
Median and interquartile ranges of the number of flowers (log scale) produced by plant clumps. SJ = *S. rubra* ssp. *jonesii*, SP = *S. purpurea* var. *montana*. Clumps that produced no flowers were excluded.

### Flower stage and pollen viability

A total of 520 flowers were staged (Table S1) weekly between 2019 and 2023.

Flower opening and pollen shedding in the two species overlapped at all sites and began earlier in *S. purpurea* var*. montana* (Table 3). Across sampling seasons which began on different dates, MB plants consistently produced flowers earlier than those at other sites, and those flowers opened and began shedding pollen earlier (Figs 3-6).

**Fig 3.**
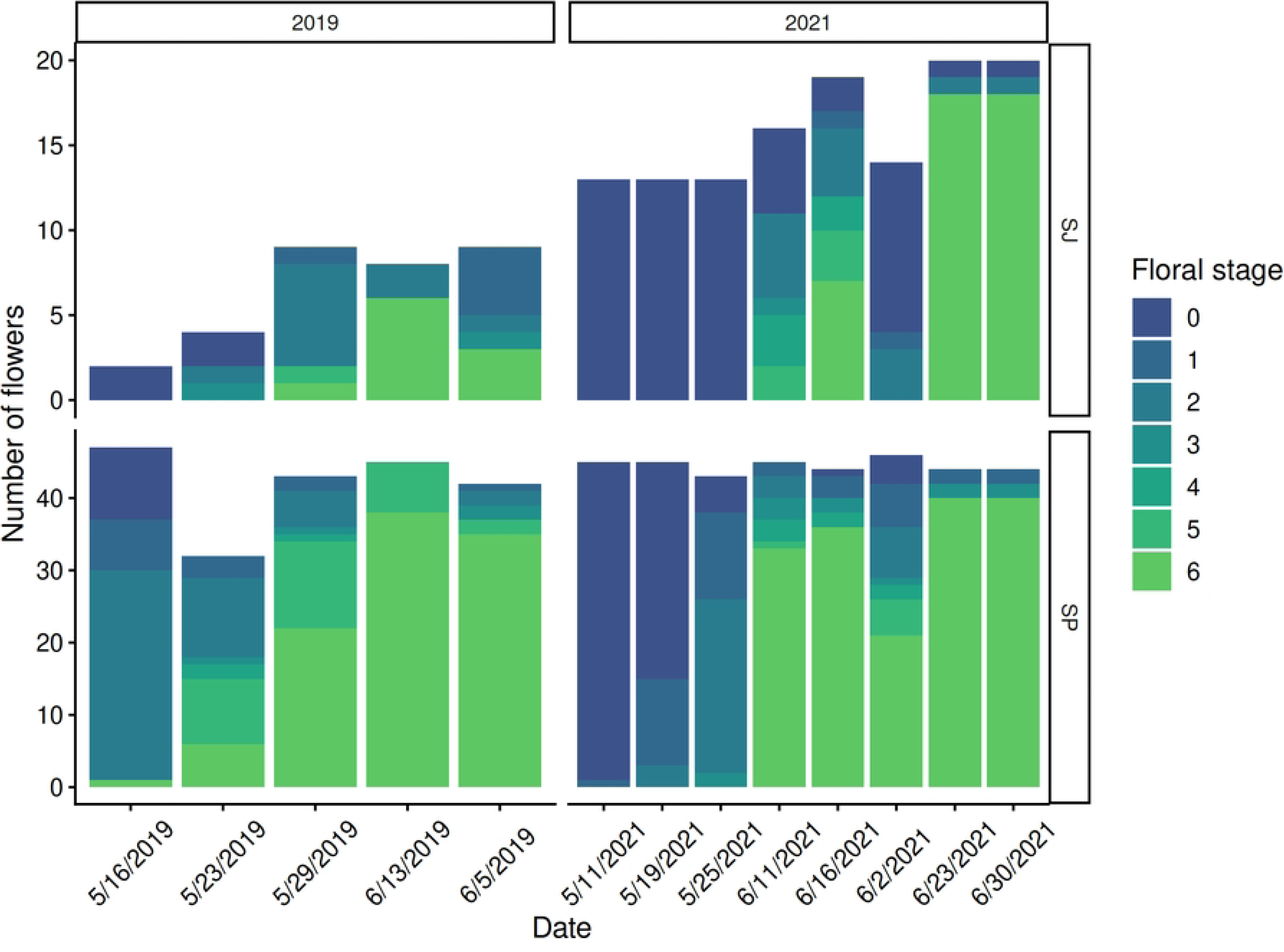
Number of flowers in each floral development stage (**Table 2**) at CM in 2019 and 2021.

**Table 3.** Results of likelihood ratio tests comparing the proportion of flowers in each stage over time and between species.

| Site | Year | Species | Date |
| --- | --- | --- | --- |
| CM | 2019 | $X^2_1 = 10.18, p = 0.001$ | $X^2_1 = 237.22, p < 0.0001$ |
| | 2021 | $X^2_1 = 26.54, p < 0.0001$ | $X^2_1 = 623.16, p < 0.0001$ |
| MB | 2019 | $X^2_1 = 107.78, p < 0.0001$ | $X^2_1 = 547.30, p < 0.0001$ |
| | 2021 | $X^2_1 = 107.48, p < 0.0001$ | $X^2_1 = 942.25, p < 0.0001$ |
| RL | 2021 | $X^2_1 = 47.06, p < 0.0001$ | $X^2_1 = 882.51, p < 0.0001$ |
| | 2022 | $X^2_1 = 43.02, p < 0.0001$ | $X^2_1 = 948.54, p < 0.0001$ |
| SV | 2023 | $X^2_1 = 63.92, p < 0.0001$ | $X^2_1 = 92.75, p < 0.0001$ |

Likelihood ratios approximately fit a $X^2$ distribution with 1 degree of freedom.

Flowering at RL occurred later than at the other three sites (Figs 3-6). Flowering at CM and SV, sites located within 3 km of each other, was temporally intermediate. Although the timing of flower opening differed among sites, some flowers of both species retained anthers throughout the sampling period at many sites. This is likely because both species continue to produce flowers throughout the season and flowers were added throughout sampling periods to reach three per clump.

At CM, we observed some open *S. purpurea* var*. montana* flowers on the first sampling days in 2019 and 2021 but did not observe open *S. rubra* ssp. *jonesii* flowers until one and three weeks later, respectively (Fig 3). *S. purpurea* var*. montana* began shedding pollen one week earlier than *S. rubra* ssp. *jonesii* in 2019 and three weeks earlier in 2021. At MB, flowers of both species were open and shedding pollen on the first sampling day of 2019 (Fig 4). In 2021, many *S. purpurea* var*. montana* flowers were already shedding pollen (≥ stage 2) on our first day of sampling, but *S. rubra* ssp. *jonesii* flowers did not begin shedding pollen until two weeks later. At RL, the first *S. rubra* ssp. *jonesii* flowers opened two weeks after *S. purpurea* var*. montana* flowers were first observed open (Fig 5). In 2022, we did not begin sampling at RL until late May, when some flowers of both species were already open and *S. purpurea* var*. montana* flowers were shedding pollen. *S. rubra* ssp. *jonesii* flowers began shedding pollen a week later. Flowers at SV followed a similar pattern to those at CM. Some *S. purpurea* var*. montana* flowers were open and shedding pollen by the first day of monitoring in 2023, and *S. rubra* ssp. *jonesii* flowers began shedding pollen two weeks later (Fig 6).

**Fig 4.**
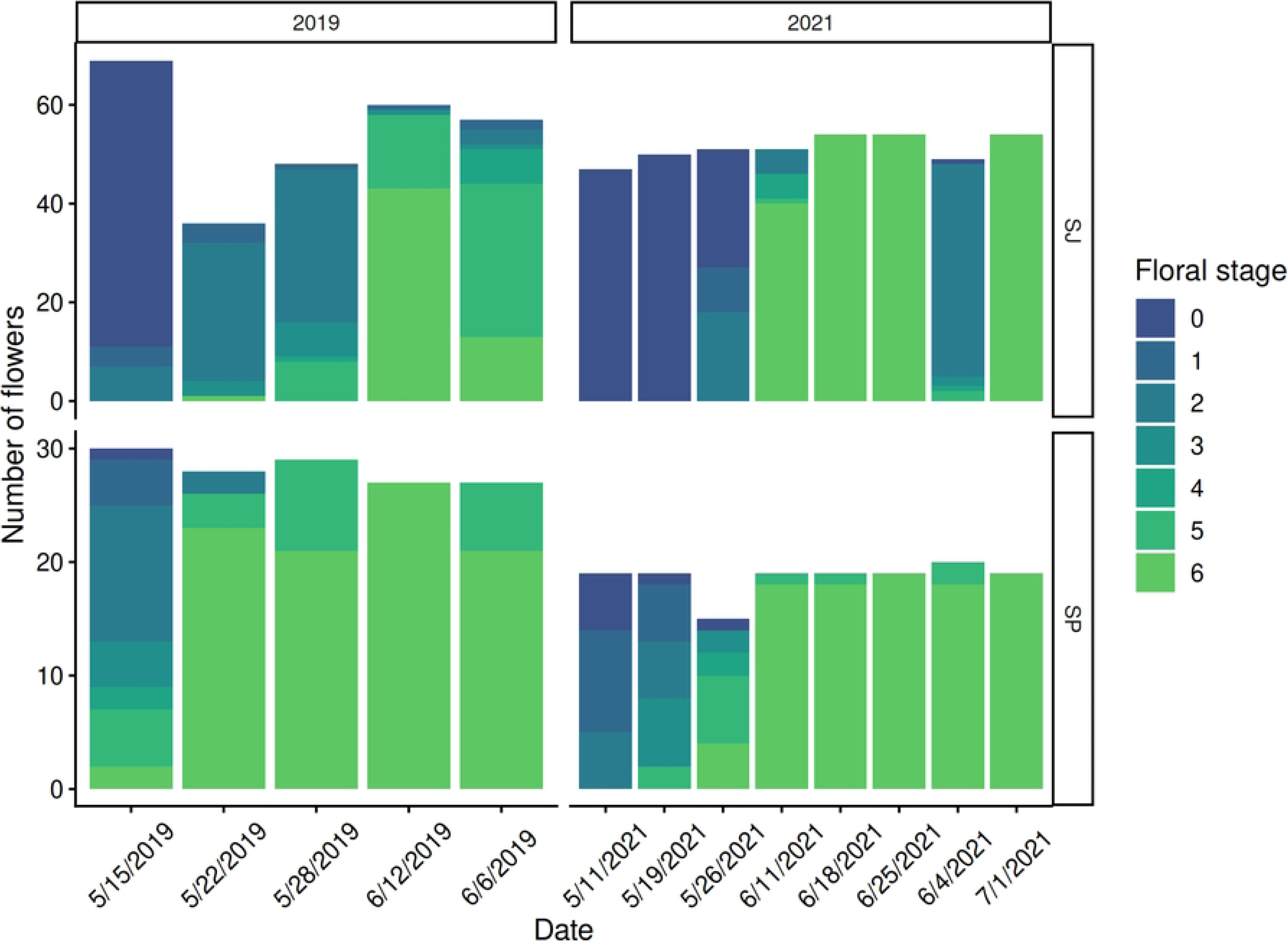
Number of flowers in each floral development stage (**Table 2**) at MB in 2019 and 2021.

**Fig 5.**
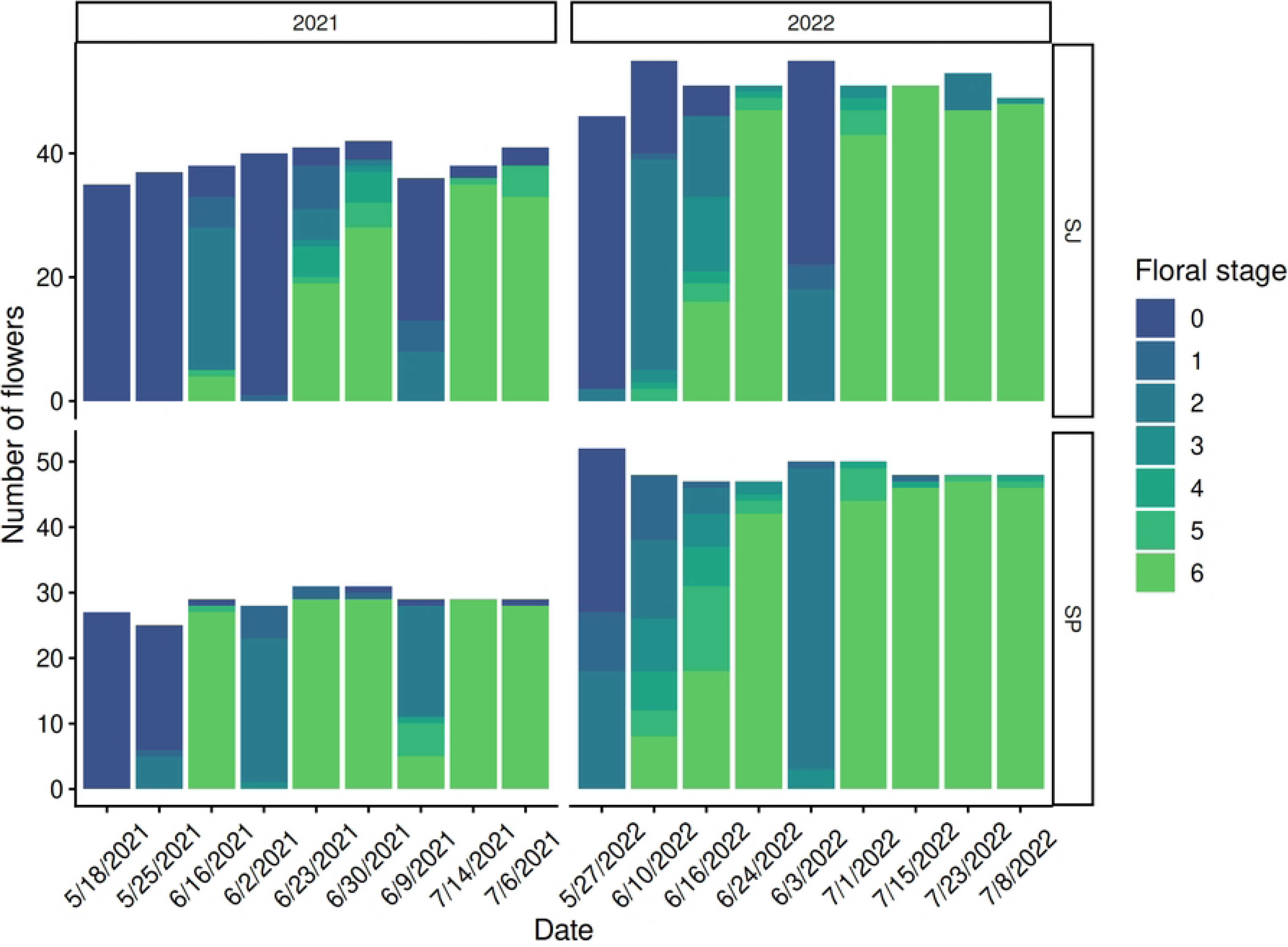
Number of flowers in each floral development stage (**Table 2**) at RL in 2021 and 2022.

**Fig 6.**
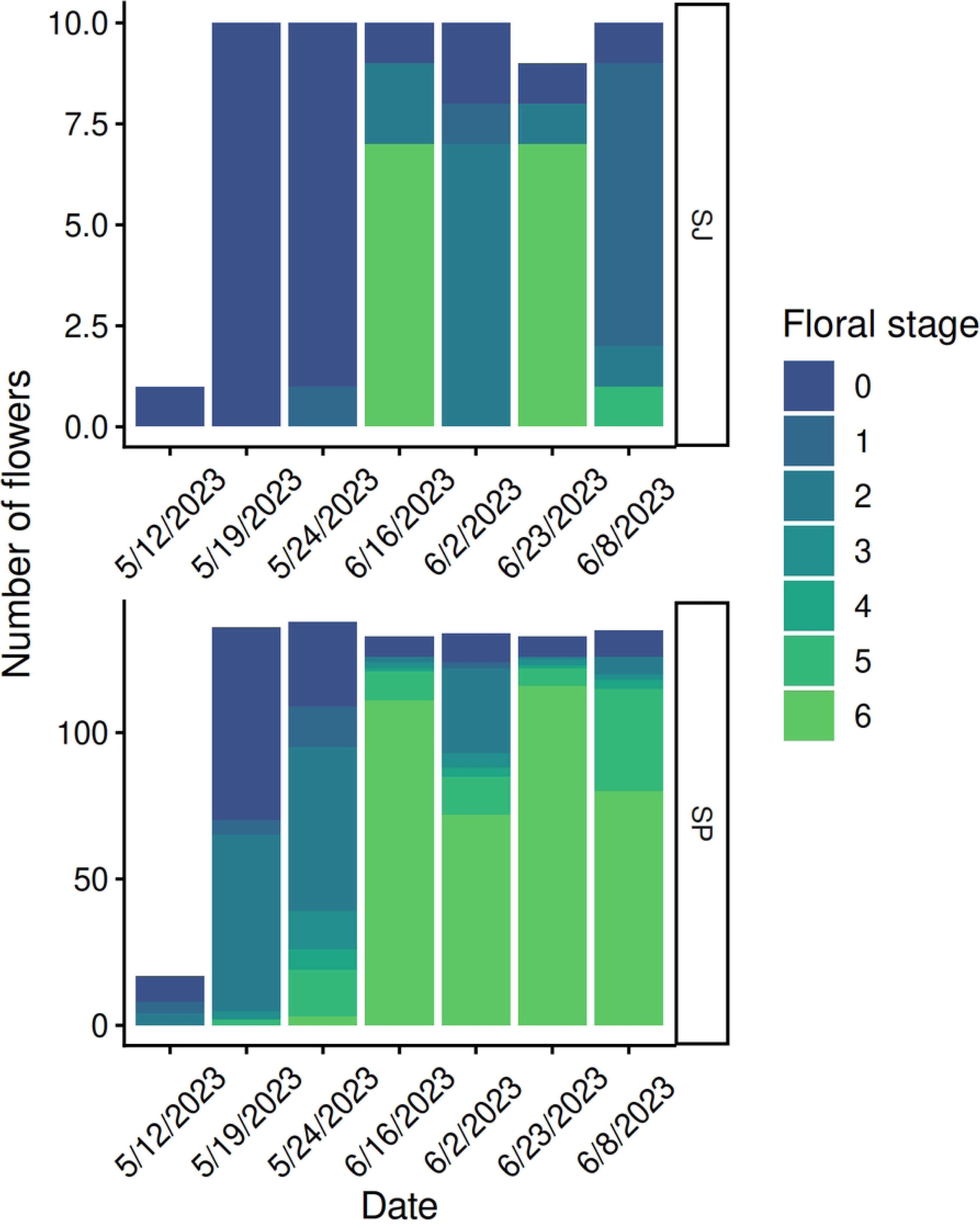
Number of flowers in each floral development stage (**Table 2**) at SV in 2023.

Of the flowers staged each week, a total of 203 *S. rubra* ssp. *jonesii* flowers and 210 *S. purpurea* var. *montana* flowers were sampled for pollen viability (S1 Table). In both species, viable pollen per anther declined as flowers advanced through stages (Spearman rank correlation; *S. rubra* ssp. *jonesii*: S = 33515203, N = 549, rho = -0.22, p < 0.0001; *S. purpurea* var*. montana*: S = 160285910, N =889, rho = -0.37, p < 0.0001), likely because pollen was shed as anthers fully dehisced. However, viable pollen per anther remained high through all stages (Fig 7). In *S. rubra* ssp. *jonesii*, viable pollen dropped from 94.5% to 93.1% between stages 1 and 6, but had the highest mean viability in stage 5 (96.7%). In *S. purpurea* var. *montana*, viable pollen dropped from its highest of 97.3% in stage 1, to its lowest of 88.5% in stage 6. Stage 6 anthers were often resting on the upward-facing surface of the style and were available for collection.

**Fig 7.**
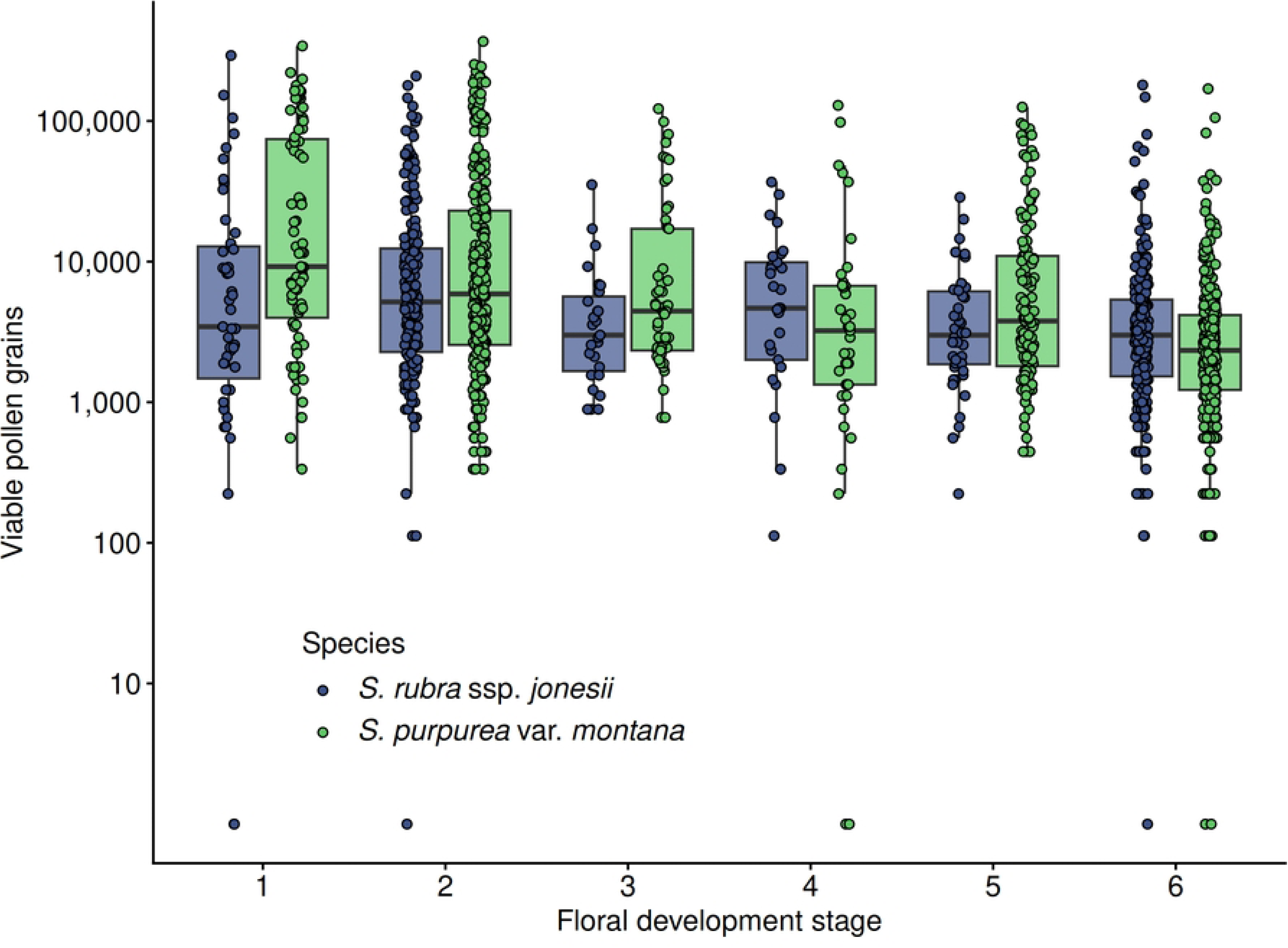
Median and interquartile ranges of the number of viable pollen grains per anther (log scale), as a function of floral development stage (**Table 2**). A total of 413 flowers were sampled for pollen viability across sites and years. Since flowers often developed through multiple stages between weekly samplings, not all flowers were sampled in each stage.

To compare viable pollen between species and among sites, we evaluated only stage 2 flowers, with mature but non-dehiscent anthers. Viable pollen differences between species varied significantly among sites (Scheirer Ray Hare test; site*species interaction: H = 13.33, df = 3, p = 0.004; Fig S1). Mann-Whitney U tests revealed significantly more viable pollen in *S. purpurea* var. *montana* than *S. rubra* ssp. *jonesii* anthers at RL (W =3268, N = 197, p = 0.0001), but no differences among species at CM, MB, or SV (CM: W = 562.5, N = 70, p = 0.13; MB: W = 226.5, N = 63, p = 0.51; SV: W = 833, N = 137, p = 0.27).

### Seed production

The proportion of *S. purpure*a var*. montana* flowers producing seeds ranged from 0.07 at CM to 1.0 at both MB and SF (Fig 8). The proportion of *S. rubra* ssp. *jonesii* flowers producing seeds ranged from 0.33 at SV to 1.0 at MB. There was a significant difference among sites in the log odds of flowers producing seeds (likelihood ratio test; site: X^2^_1_ = 16.41, p = 0.012) but no difference between species and no site x species interaction (species: X^2^ = 0.0001, p = 0.99, interaction: X^2^ = 5.55, p = 0.24). To determine site differences, we evaluated the 95% CIs around each site estimate. The proportion of flowers of both species that produced seeds at CM was significantly lower than the proportions at MB, RL, SF, or SV.

**Fig 8.**
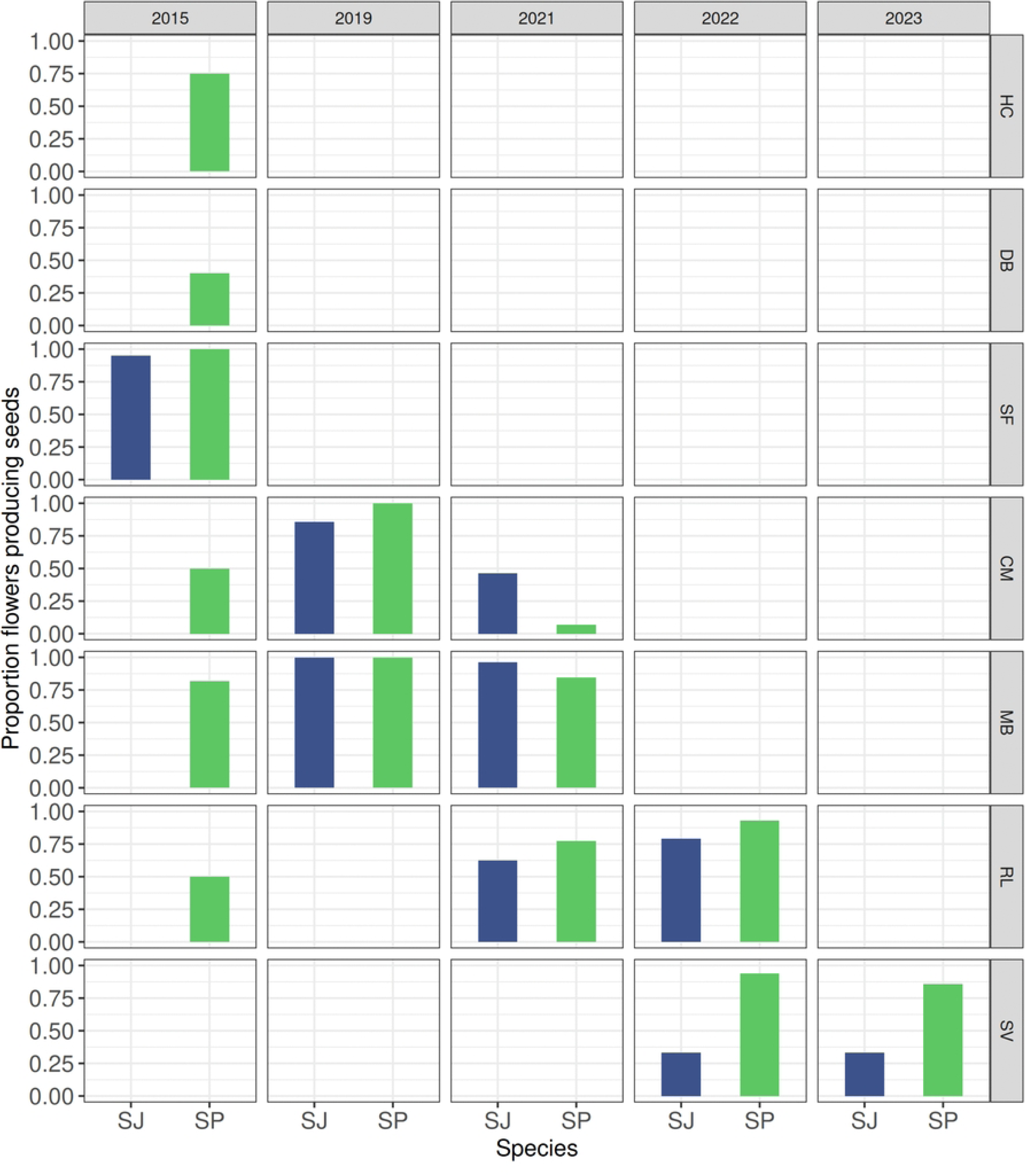
Proportion of flowers producing seeds at seven sites between 2015 and 2023. In 2015, only *S. purpurea* var. *montana* flowers were examined at four of the five sites studied. Thus, there are more years of data for *S. purpurea* var. *montana* (SP) than for *S. rubra* ssp. *jonesii* (SJ).

Among flowers that produced seeds, the range of seeds per flower was highly variable, with at least one flower at each site producing only one seed (Fig 9). The difference in mean seed count between species differed significantly among sites (Scheirer Ray Hare test; site*species interaction: H = 18.16, df = 3, p = 0.0004; site: H=10.67, df=3, p=0.014; species: H=22.17, df=1, p<0.001). Mann-Whitney U tests revealed that *S. purpurea* var. *montana* produced more seeds than *S. rubra* ssp. *jonesii* at all sites except MB (CM: W = 34.5, N = 30, p = 0.002; MB: W = 834.5, N = 85, p = 0.23; RL: W = 1033, N = 121, p < 0.001; SV: W = 58, N = 144, p = 0.007).

**Fig 9.**
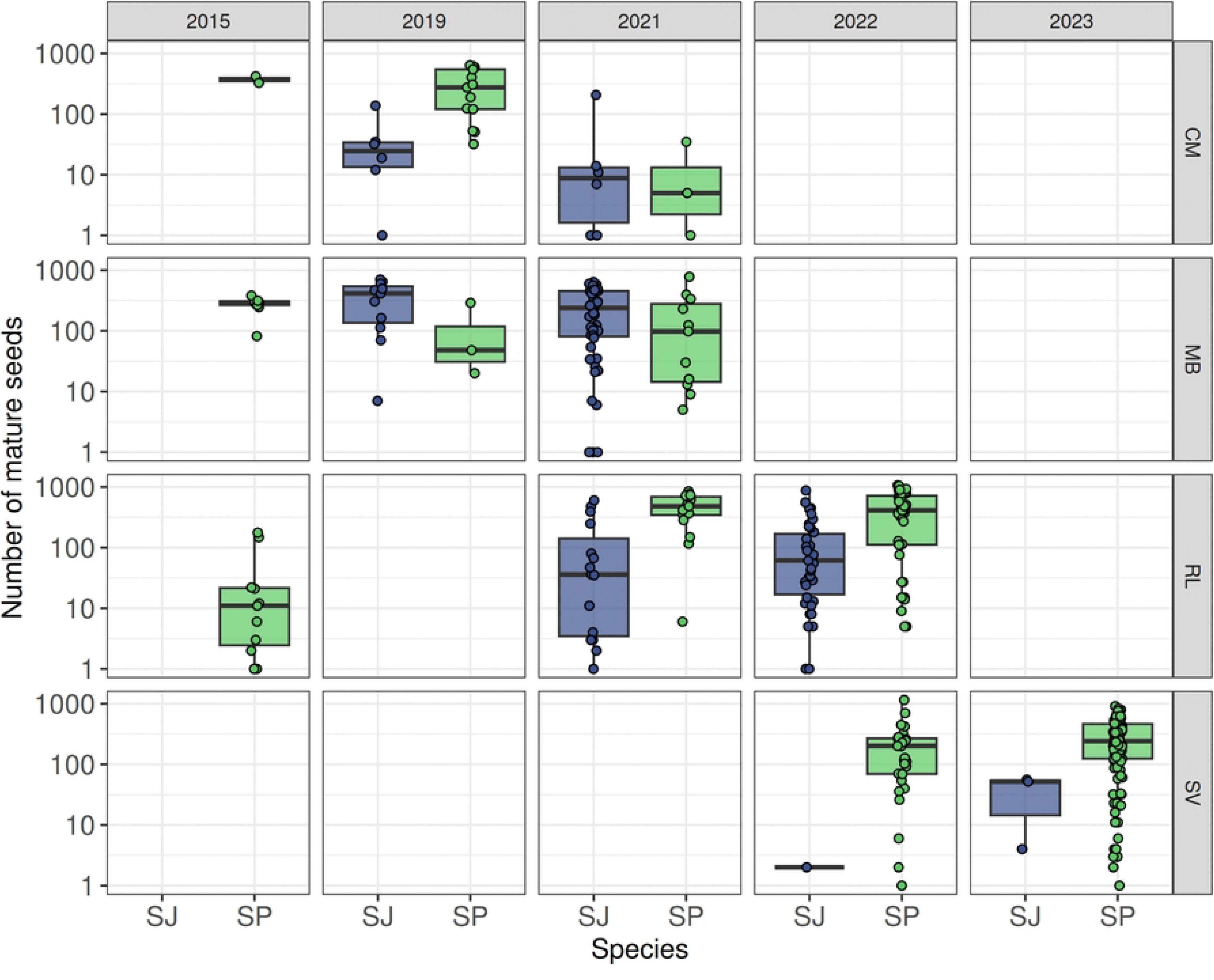
Median and interquartile ranges of the number of seeds (log scale) produced by flowers. SJ = *S. rubra* ssp. *jonesii*, SP = *S. purpurea* var. *montana*. Flowers that produced no seeds were excluded.

### Seed size and germination

Germination success of seeds at RL ranged from 0 to 60% in *S. rubra* ssp. *jonesii* and from 0 to 100% in *S. purpurea* var*. montana* across samples. Seeds of *S. purpurea* var*. montana* had significantly greater mass (likelihood ratio test, X^2^ = 15.31, p < 0.001) and were significantly more likely to germinate (X^2^ = 17.24, p < 0.001) than those of *S. rubra* ssp. *jonesii*. However, when these two variables were evaluated in the same model, there was no difference between species independent of the effect of mass (mass: X^2^ = 5.39, p = 0.02; species: X^2^ = 1.42, p = 0.23; Fig 10).

**Fig 10.**
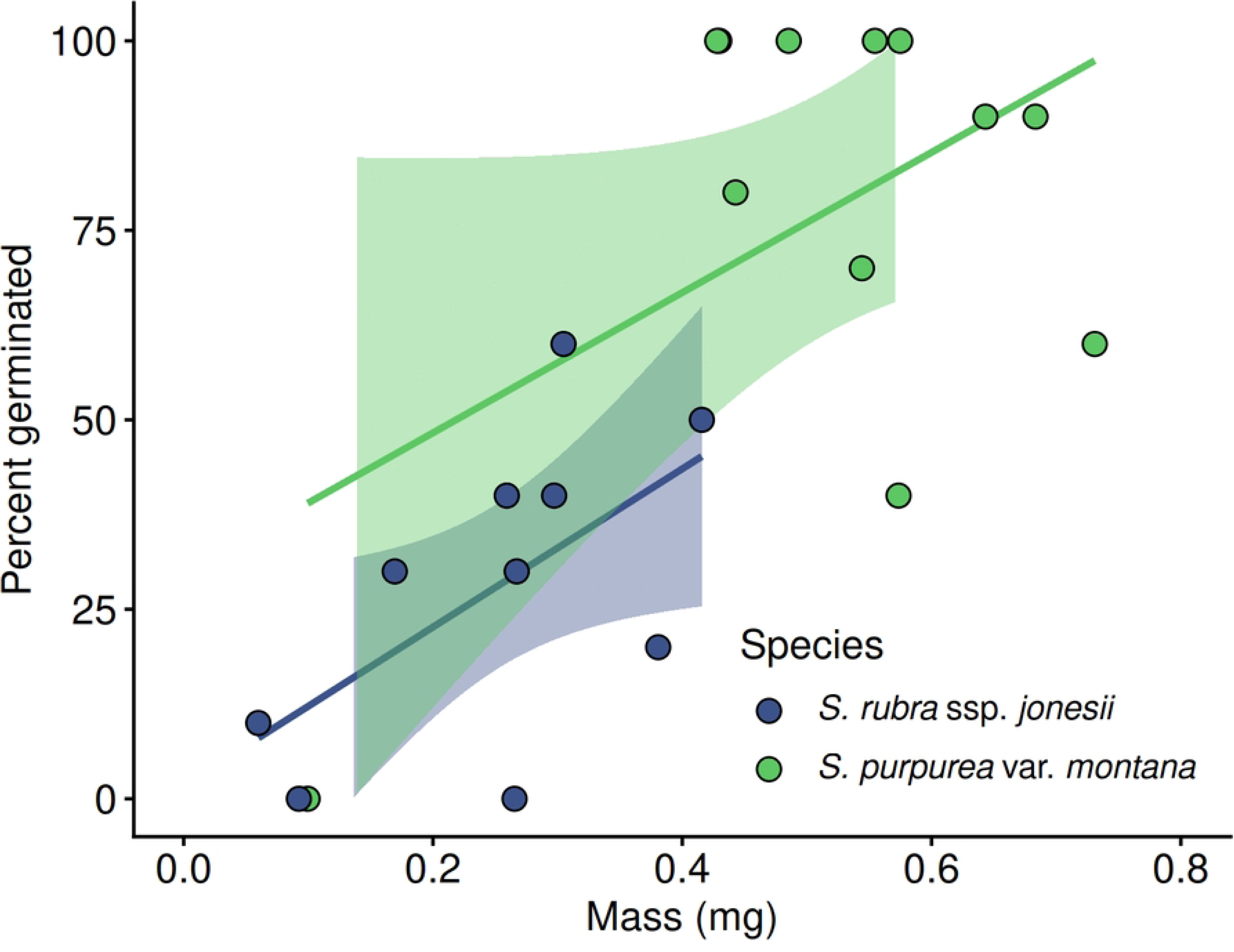
Percent of seeds that germinated as a function of seed mass. Each point represents one flower, and seed mass was averaged across 5 seeds. Ten different seeds were used to measure germination success.

## Discussion

### Reproductive Effort

This study quantified considerable variation in reproductive effort between imperiled *Sarracenia* species and among sites. While species did not differ in their probability of flowering, *S. jonesii* generally produced more flowers per clump. Using management convention, we defined clumps as groups of interconnected stems, assumed to constitute one ramet (44) and including one or more rosettes. The number of flowers per clump may be a function of clump size, which has no standard measure and so was not quantified. Therefore, our data do not provide conclusive evidence for higher energetic investment in flower production by *S. rubra* ssp. *jonesii* plants.

Pollen production per anther also varied by species, but these patterns were not consistent among sites. *S. rubra* ssp. *jonesii* produced more pollen than *S. purpurea* var. *montana* at one location and less in another, and at two sites pollen production did not differ significantly. Thus, when flower and pollen production are considered together, we saw no evidence of intertaxonomic differences in reproductive effort. This study also found no evidence of tradeoffs at the taxon level between flower abundance and pollen per flower. Because ovules were not enumerated in this study, their potential contribution to differences in reproductive effort are not known. However, data from Ne’eman et al. (25) showed little evidence that pollen or ovules limit seed production in *Sarracenia purpurea*.

### Phenology

The absolute timing of flower production and progression through floral development stages differed among sites, likely due to differences in temperature related to elevation (54) and in light availability due to canopy cover (55, 56). *S. purpurea* var*. montana* flowered earlier each season than *S. rubra* ssp. *jonesii* at all sites, but species displayed considerable temporal overlap in production of pollen-shedding flowers. As individual flowers aged, they contained less viable pollen, due to both pollen loss and stamen drop. The gradual shedding of stamens could also indicate reduced opportunities for pollinator-mediated outcrossing or hybridization as flowers age. Little is known about the visitation rates of pollinators over the age of a flower, nor about the phenology of nectar rewards, but the high viability of pollen (≥ 88% throughout a flower’s lifespan) suggests that outcrossing via pollen dispersal remains possible even when all stamens have been shed.

High pollen viability even among stage 6 flowers, in which all stamens have been shed, raises the possibility of self-pollination. Once shed, anthers containing viable pollen fall on the stigma, which hangs beneath the ovary on which stamen are attached. Self-pollination occurs in this genus (30) and could provide fertilization assurance if flowers do not receive outcrossed pollen via animal visitors, as is seen in many angiosperms (57). If so, we would expect stigma to be receptive before anthers begin shedding pollen to aid outcrossing, and to remain receptive through anther shedding to facilitate selfing. While no dichogamy has been observed in this genus (28), the phenology of stigma receptivity in these species is unknown. In addition, although at least one taxon (*S. purpurea*) is capable of self-fertilization (25), neither species’ rates of self-fertilization on the landscape is known. This pre-zygotic phenomenon, along with actual mating patterns and how floral visitors might affect hybridization (e.g., 58), warrant further study.

### Reproductive Output

Although we observed no consistent patterns in reproductive effort between species, strong patterns in reproductive output emerged. Absolute seed production differed among sites, potentially due to differences in light availability (56) or conspecific density (55) . However, over all sites, *S. purpurea* var. *montana* flowers produced more seeds per ovary than *S. rubra* spp. *jonesii*. This could be due to differences in ovule production or pollination success. *S. purpurea* var. *montana* flowers also produced seeds of greater mass, consistent with a previous study (52) and displaying no evidence of a tradeoff between seed number and size (59, 60). In addition, for the single population examined, *S. purpurea* var. *montana* seeds exhibited higher germination rates. These differences, together, suggest that *S. purpurea* var. *montana* plants invest more into seed production and quality than do *S. rubra* ssp. *jonesii* plants. Although larger seeds do not always have higher germination rates (61), these species’ investment in mass is correlated with greater fitness through the germinant phase of offspring development.

Interspecific differences in reproductive output could be the result of differences in total energy budgets or in energy allocation. Both species receive nutrients from decaying animal material in their pitchers (62, 63, 64), yet they acquire those nutrients through different processes. As in other *Sarracenia*, *S. rubra* ssp. *jonesii* pitchers likely digest prey items with endogenously-produced enzymes (65, 66, 67). In contrast, *S. purpurea* var. *montana* pitchers acquire nutrients through the waste products of the inquiline community inside their pitchers, members of which digest and break down prey (68, 69). As a result, *S. purpurea* var. *montana* need not continually secrete digestive enzymes into pitchers (70). This method of acquiring nutrients might cost plants less, release more nutrients, and do so more consistently over the growing season. This could give *S. purpurea* var. *montana* a larger energy budget than in taxa like *S. rubra* ssp. *jonesii* that lack inquiline communities. Prey and inquiline manipulation studies (e.g., 71, 72, 73) that quantify the relative sensitivity of these two species’ growth and reproductive effort to variation in prey capture could reveal differences in these species’ energy budgets and how they are allocated. Further, as nutrients are acquired, *S. purpurea* var. *montana* might allocate them differently than *S. rubra* ssp. *jonesii*. Isotope studies that examine how nutrients from prey are allocated to vegetative versus reproductive tissues could confirm the energetic differences underlying interspecific trends in reproduction.

### Hybridization Potential

Despite *S. purpurea* var. *montana* consistently flowering earlier, abundant viable pollen from both species was available throughout the reproductive season. This, along with the movement of individual floral visitors among taxa (Kennedy, pers. obs.), could introduce opportunities for interspecific hybridization. Although scapes are different heights, this vertical stratification, which results in pollinator segregation in some cases (74), provides a barrier to cross-pollination that is porous at best. Phenotypic hybrids between these two species have been observed at three of the sites where taxa co-occur (40, 49, 75, 76). While hybrid individuals were excluded from analysis, their presence and relatively high fitness could lead to the loss of species identity, further complicating conservation efforts for these listed taxa (54, 77). Alternatively, if reproduction in these taxa is primarily asexual, emergence of novel hybrids with relatively high pollen and seed production might not be a challenge to the parental taxa (78). For now, though, potential hybrid incompatibility (79) or niche differentiation (80) remains uncharacterized at these sites.

### Future Directions

This study identified inconsistent differences in male reproductive effort between taxa. In a study of a single site (SF), Uzzell (81) reported that ovule production by *S. rubra* ssp. *jonesii* and *S. purpurea* var. *montana* was statistically indistinguishable, but that phenotypic hybrids made more ovules than *S. rubra* ssp. *jonesii*. Overall differences in ovule production among sites and species remain unknown, however. We showed that *S. purpurea* var. *montana* produced more seeds per ovary than its more imperiled congener at sites where the taxa co-occur. This could lead to greater dominance of *S. purpurea* var. *montana* on the landscape, if seed production is more important than clonal growth for population growth. In contrast, if clonal growth is a greater driver of population growth, *S. rubra* ssp. *jonesii*, whose clumps are larger and contain more pitchers, could crowd suitable sites and become dominant on the landscape. However, the relative importance of vegetative versus sexual reproduction remains to be explored in these species and without high resolution genetic markers, accurately distinguishing between ramets and genets is impossible. Further, the genetic relatedness of adjacent rosettes is not known. These could be clonal or could be close relatives if seeds, whose mean dispersal distance is on the order of centimeters (82), germinate near maternal plants.

Although our laboratory experiments showed relatively high rates of germination, we have never observed a *Sarracenia* seedling at any of these field sites. Therefore, the steps between seed production, seed dispersal, germination, and successful establishment under field conditions could be a bottleneck preventing taxa from showing more genetic diversity or establishing new populations. The sites to which these gravity- and water-dispersed seeds move might be important, as germination could be light-limited and the chances of reaching inhospitable environments is high. In addition, small germinants might be displaced by seasonal flooding, a late summer phenomenon at each of these sites (Rhode Ward, pers. obs.). Understanding seed fates is essential for successful management of these populations.

Although phenological overlap in pollen viability could allow interspecific hybridization, the phenology of stigma receptivity is unknown. This could limit opportunities for both hybridization and intraspecific outcrossing, and measuring this could give insight into outbreeding potential. While this and previous studies (25) seem to indicate that these species are not pollen limited, they might be pollinator limited. Both species rely on a suite of generalist pollinators, which might also be shared with other community members whose flowers are receptive at the same times; this could increase (facilitation) or decrease (competition) *Sarracenia* pollination (83).

The genetic identity of embryos is also unknown. In sites where these taxa co-occur, significant overlap in pollen production, combined with a shared suite of generalist pollinators, could allow hybridization. The genetic identity of mature phenotypic hybrids observed on the landscape is also unclear, but these individuals produce pollen and seed numbers similar to those reported here (81). Hybrids’ phenological overlap with parental taxa (81) could also prove problematic (e.g. 84). Hybrid fitness on the landscape, the ability of hybrids to compete with parental taxa for limited resources like light and space, and the ways in which demography of the two taxa might interact (e.g., Berry and Cleavitt 2021) is also unknown. Resolving each of these questions could be key to the continued survival of both imperiled groups.

## Acknowledgements

We thank Kathy Reimer, Highlands Biological Station (James Costa and Jason Love), The Nature Conservancy, and the North Carolina Forest Service (Michael Santucci and Jordan Luff) for site access.

---

**S1 Fig.**
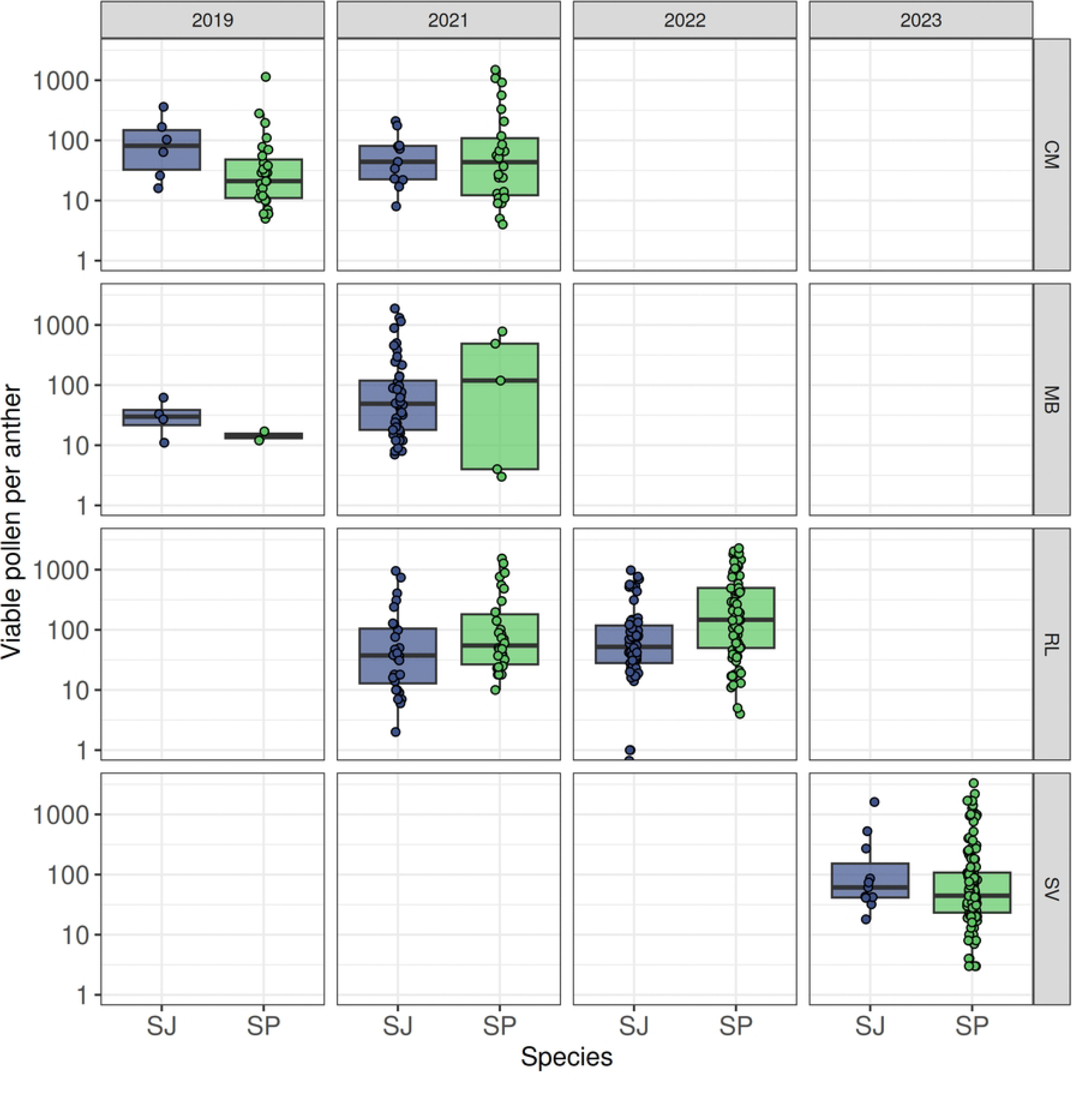
Number of viable pollen grains per anther on stage 2 flowers. SJ = *S. rubra* ssp. *jonesii*; SP = *S. purpurea* var. *montana*.

**S1 Table.** Number of flowers sampled for stage, pollen viability, and seed production.

| Site | Year | Species | Clumps | Flowers sampled for: |  |  |
| --- | --- | --- | --- | --- | --- | --- |
|  |  |  |  | Stage | Pollen | Seeds |
| DB | 2015 | SP | 8 |  |  | 9 |
| HC | 2015 | SP | 4 |  |  | 4 |
| CM | 2015 | SP | 4 |  |  | 4 |
| CM | 2019 | SJ | 5 | 9 | 6 | 7 |
| CM | 2019 | SP | 21 | 54 | 34 | 13 |
| CM | 2021 | SJ | 9 | 20 | 14 | 13 |
| CM | 2021 | SP | 24 | 50 | 45 | 43 |
| MB | 2015 | SP | 8 |  |  | 9 |
| MB | 2019 | SJ | 26 | 76 | 36 | 11 |
| MB | 2019 | SP | 10 | 31 | 19 | 3 |
| MB | 2021 | SJ | 20 | 54 | 52 | 53 |
| MB | 2021 | SP | 12 | 21 | 17 | 13 |
| RL | 2015 | SP | 21 |  |  | 19 |
| RL | 2021 | SJ | 19 | 43 | 39 | 3 |
| RL | 2021 | SP | 26 | 34 | 34 | 3 |
| RL | 2022 | SJ | 26 | 59 | 53 | 3 |
| RL | 2022 | SP | 28 | 59 | 56 | 3 |
| SF | 2015 | SJ | 20 |  |  | 1 |
| SF | 2015 | SP | 11 |  |  | 1 |
| SV | 2022 | SJ | 1 |  |  | 3 |
| SV | 2022 | SP | 20 |  |  | 5 |
| SV | 2023 | SJ | 4 | 4 | 3 | 3 |
| SV | 2023 | SP | 64 | 6 | 5 | 6 |

## Notes

### Competing Interest Statement

The authors have declared no competing interest.

## References

1. Stearns S. The Evolution of Life Histories. Oxford University Press; 1992.

2. Bazzaz F, Ackerly D, Reekie E. Reproductive Allocation in Plants. In: Seeds: The Ecology of Regeneration in Plant Communities. 2nd ed. 2000.

3. Hirshfield M, Tinkle D. Natural selection and the evolution of reproductive effort. Proceedings of the National Academy of Science. 1975;76(2):2227–2231. doi:10.1073/pnas.72.6.2227

4. Thompson K, Stewart A. The measurement and meaning of reproductive effort in plants. The American Naturalist. 1981;117(2):205–211. doi:https://www.jstor.org/stable/2460502

5. Karlsson P, Méndez M. The Resource Economy of Plant Reproduction. In: Reproductive Allocation in Plants. 2005.

6. Lloyd D. Sexual strategies in plants I. An hypothesis of serial adjustment of maternal allocation during one reproductive session. New Phytologist. 1980;86:69–79. 10.1111/j.1469-8137.1980.tb00780.x

7. Casper B, Niesenbaum R. Pollen versus resource limitation of seed production: a reconsideration. Current Science. 1993;65(3):210–214.

8. Aizen M, Harder L. Expanding the limits of the pollen-limitation concept: effects of pollen quantity and quality. Ecology. 88(2):271–281. 10.1890/06-1017

9. Gomez J, Abdelaziz M, Lorite J, Munoz-Pajares AJ, Perfectti F. Changes in pollinator fauna cause spatial variation in pollen limitation. Journal of Ecology. 2010;98:1243–1252. 10.1111/j.1365-2745.2010.01691.x

10. Weiner J, Campbell L, Pino J, Echarte L. The allometry of reproduction within plant populations. Journal of Ecology. 2009;97:1220–1233. 10.1111/j.1365-2745.2009.01559.x

11. Dorken M, van Kleunan M, Stift M. Costs of reproduction in flowering plants. New Phytologist. 2025;247:55–70. 10.1111/nph.70166

12. Barrett S. Understanding plant reproductive diversity. Philosophical Transactions of the Royal Society B. 2010;365(1537):99–109. 10.1098/rstb.2009.0199

13. Zhou Y, Li X, Zhao Y, et al. Divergences in reproductive strategy explain the distribution ranges of *Vallisneria* species in China. Aquatic Botany. 2016;132:41–48. 10.1016/j.aquabot.2016.04.005

14. Mochizuki J, Itagaki T, Blue Y, Ito M, Sakai S. Ovule and seed production patterns in relation to flower size variations in actinomorphic and zygomorphic flower species. AoB Plants. 2019;11(5):plz061. 10.1093/aobpla/plz061

15. Bawa K, Ingty T, Revell L, Shivaprakash K. Correlated evolution of flower size and seed number in flowering plants (monocotyledons). Annals of Botany. 2018;123(1). doi:doi:%2010.1093/aob/mcy154

16. Sun HQ, Huang BQ, Yu XH, Tian CB, Peng XX, An DJ. Pollen limitation, reproductive success and flowering frequency in single-flowered plants. Journal of Ecology. 2018;106:19–30. 10.1111/1365-2745.12834

17. Kang X, Liu Y, Wu X, et al. Alpine grassland community productivity and diversity differences influence significantly plant sexual reproduction strategies. Proceedings of the National Academy of Science Nexus. 2024;3(8). doi:doi:%2010.1093/pnasnexus/pgae297

18. Liao D, Lei J, Wang Y, Bao Y, Zhang Z, Wang J. Sex-specific responses of sexual reproduction, clonal reproduction, and vegetative growth to environmental (biotic and abiotic) factors in the clonal dioecious plant *Acer barbinerve*. Plants. 2025;14:596. 10.3390/plants14040596

19. Christie K, Fraser L, Lowry D. The strength of reproductive isolating barriers in seed plants: Insights from studies quantifying premating and postmating reproductive barriers over the past 15 years. Evolution. 2022;76(10). doi:DOI:%2010.1111/evo.14565

20. Jiminez-Lopez F, Arista M, Tallavera M, et al. Multiple pre- and postzygotic components of reproductive isolation between two co-occurring *Lysimachia* species. New Phytologist. 2023;238:874–887. 10.1111/nph.18767

21. Lan Z, Song Z, Wang Z, et al. Antagonistic RALF peptides control an intergeneric hybridization barrier on Brassicaceae stigmas. Cell. 186:4773–4787. doi:10.1016/j.cell.2023.09.003

22. Bureš P., Šmarda P., Rotreklová O., Oberreiter M., Burešová M., Konečný J., Knoll A., Fajmon K. Šmerda J. Pollen viability and natural hybridization of Central European species of *Cirsium*. Preslia. 2010;89:3.

23. Richardson C, Gibbons L. Pocosins, Carolina bays, and mountain bogs. In: Biodiversity of the Southeastern United States. John Wiley & Sons; 1993.

24. Weakley A, Schafale M. Classification of the Natural Communities of North Carolina. 4th ed. 2024.

25. Ne’eman G, Ne’eman R, Ellison AM. Limits to reproductive success of *Sarracenia purpurea* (Sarraceniaceae). Am J Bot. 2006;93(11):1660–1666. doi:10.3732/ajb.93.11.1660

26. Meyers-Rice B. Rare Sarracenia poaching and the ICPS. Carnivorous Plant Newsletter. 2001;30(2):43–50.

27. Jennings D, Rohr J. A review of the conservation threats to carnivorous plants. Biological Conservation. 2011;144(5):1356–1363. 10.1016/j.biocon.2011.03.013

28. Burr CA. The Pollination Ecology of Sarracenia Purpurea in Cranberry Bog, Weybridge, Vermont (Addison County). M.S. Thesis. Middlebury College; 1979.

29. Thomas K, Cameron D. Pollination and fertilization in the pitcher plant (*Sarracenia purpurea* L). American Journal of Botany. 1986;73:678.

30. Sheridan P, Mills R. Genetics of anthocyanin deficiency in *Sarracenia* L. HortScience. 1998;33:1042–1045.

31. Schnell D. Carnivorous Plants of the United States and Canada. 2nd ed. Timber Press; 2002.

32. McDaniel S. The genus Sarracenia (Sarraceniaceae). Bulletin of Tall Timbers Research Station. 1971;9:1–36.

33. Vogel S. Remarkable nectaries: structure, ecology, organophyletic perspectives. II. Nectarioles. Flora. 1998;193:1–29.

34. Ellison A, Butler E, Hicks E, et al. Phylogeny and Biogeography of the Carnivorous Plant Family Sarraceniaceae. PLoS ONR. 2012;7(6):e39291. 10.1371/journal.pone.0039291

35. Mellichamp T. New Names for Natural Hybrids in Sarracenia. Carnivorous Plant Journal. 34(7):112. doi:doi.%2010.55360/cpn374.lm492

36. Baldwin E, McNair M, Leebens-Mack J. Rampant chloroplast capture in *Sarracenia* revealed by plastome phylogeny. Frontiers in Plant Science. 2023;14:1237749.

37. Sandlin I, Kaye T. Ploidy-Mediated Reproductive Barriers and Hybridization between a Rare and a Common *Castilleja* Species (Orobanchaceae). Natural Areas Journal. 46(3):199–206. 10.3375/2162-4399-46.3.4

38. Schnell D, Determann R. *Sarracenia purpurea* L. ssp. *venosa* (Raf.) Wherry var. montana Schnell & Determann (Sarraceniaceae): a new variety. Castanea. 1997; 62:60–62.

39. Mellichamp T. The Sarracenia pitcher plants and bog gardening. Sibbaldia. 6:79-99.

40. Brasseur T. Helping conservationists easily identify *Sarracenia purpurea* var. *montana*, *S. jonesii*, and their hybrids in the field. *Capstone*, The UNC Asheville Journal of Undergraduate Scholarship. 2021;34. doi:https://janeway.uncpress.org/capstone/article/id/3537/

41. Godt M, Hamrick J. Genetic Structure of Two Endangered Pitcher Plants, *Sarracenia jonesii* and *Sarracenia oreophila* (Sarraceniaceae). American Journal of Botany. 1996;83(8):1016–1023.

42. Flora of North America Editorial Committee. Sarracenia jonesii. In: Flora of North America. Vol 8. Magnoliophyta: Paeoniaceae to Ericaceae. 2009.

43. Massey J, Otte D, Atkinson T, Whetstone R, Sizemore S. An Atlas and Illustrated Guide to the Threatened and Endangered Vascular Plants of the Mountains of North Carolina and Virginia. Southeastern Forest Experiment Station; 1983.

44. Murdock N. Seabeach Amaranth Listing. Federal Registry. 1993;58:18035–18042.

45. R Foundation for Statistical Computing. R: A Language and Environment for Statistical Computing. R package version 2026 v4.6.1.

46. Wenk E, Falster D. Quantifying and understanding reproductive allocation schedules in plants. Ecology and Evolution. 2015;5(23):5521–5538. 10.1002/ece3.1802Digital Object Identifier (DOI)

47. Kearns C, Inouye D. Techniques for Pollination Biologists. University Press of Colorado; 1993.

48. Christensen R. ordinal - Regression Models for Ordinal Data. R package version 2026 v4.6.1.

49. Hillegass K. Potential for hybridization between Sarracenia species in sympatry. UNC Asheville Journal of Undergraduate Research. 2021;34(1).

50. Gay W, Parker G, Zappia M. Analyzing reproductive effort, reproductive output, and pitcher morphology variation to monitor *Sarracenia purpurea* var. *montana* Populations in Southern Appalachia. UNC Asheville Journal of Undergraduate Research. 2024;37.

51. Mangiafico S. *rcompanion*: Functions to Support Extension Education Program Evaluation. R package version 2026 v4.6.1.

52. Ellison A. Interspecific and intraspecific variation in seed size and germination requirements of *Sarracenia* (Sarraceniaceae). American Journal of Botany. 2001;88(3):429–437.

53. Bates D, Machler M, Bolker B, Walker S. Fitting Linear Mixed-Effects Models Using lme4. 2015;67(1):1–48. doi:doi:10.18637/jss.v067.i01

54. Chang N, Eserman L, Carmichael A, et al. Ecological niche modeling reveals habitat differentiation and climatic vulnerability in two imperiled, sympatric southern Appalachian carnivorous plants. American Journal of Botany. 2026;113(5):e70194. 10.1002/ajb2.70194

55. Brewer JS. Inter- and intraspecific competition and shade avoidance in the carnivorous pale pitcher plant in a nutrient-poor savanna. American Journal of Botany. 2019;106(1):81–89. doi:https://www.jstor.org/stable/26617191

56. Segala M, Horner J. The effects of light availability, prey capture, and their interaction on pitcher plant morphology. Plant Ecology. 224:539–548. 10.1007/s11258-023-01320-6

57. Busch J, Delph L. The relative importance of reproductive assurance and automatic selection as hypotheses for the evolution of self-fertilization. Annals of Botany. 2011;109(3):553–562.

58. Estévez Manso Galán L, Antonetti M, Ibañez AC, Sérsic AN, Cocucci AA. Phenotypic selection patterns in a hybrid zone between two *Calceolaria* species with contrasting pollinators: insights from field surveys and fitness assessments. New Phytologist. 2024; 243:440–450. doi:10.1111/nph.19775

59. Lazaro A, Larrinaga A. A multi-level test of the seed number/size trade-off in two Scandinavian communities. PLoS One. 2018;13(7): e0201175. 10.1371/journal.pone.0201175

60. Qui T, Andrus R, Aravena M, et al. Limits to reproduction and seed size-number trade-offs that shape forest dominance and future recovery. Nature Communications. 2022;13:2381.

61. Maleki K, Vandelook F, Saatkamp A, Maleki K, Heshmati S, Soltani. Global Patterns in the Evolutionary Relations Between Seed Mass and Germination Traits. Ecology and Evolution. 2025;15:e71543. 10.1002/ece3.71543

62. Adamec L. Mineral nutrition of carnivorous plants: A review. The Botanical Review. 1997;63:273–299.

63. Ellison A, Gotelli N. Nitrogen availability alters the expression of carnivory in the northern pitcher plant, Sarracenia purpurea. Proceedings of the National Academies of Science. 2002;99(7):4409–4412. 10.1073/pnas.022057199

64. Karagatzides J, Butler J, Ellison A. The pitcher plant Sarracenia purpurea can directly acquire organic nitrogen and short-circuit the inorganic nitrogen cycle. PLoS One. 2009;4(7). doi:DOI:%2010.1371/journal.pone.0006164

65. Brandon A. A Comparison Of The Enzyme Profiles Of Two Pitcher Plant Species (Sarracenia jonesii and Sarracenia purpurea var. montana) And Their Hybrids. *Capstone*, The UNC Asheville Journal of Undergraduate Scholarship. 2019;32(1). doi:https://janeway.uncpress.org/capstone/article/id/3268/

66. Koller-Peroutka M, Krammer S, Pavlik A, Edlinger M, Lang I, Adlassnig W. Endocytosis and Digestion in Carnivorous Pitcher Plants of the Family Sarraceniaceae. Plants (Basel). 2019;8(10). doi:doi:%2010.3390/plants8100367

67. Eilenberg H, Zilberstein A. Carnivorous pitcher plants: towards understanding the molecular basis of prey digestion. In: Vol 5. 1st ed. 2008.

68. Bradshaw W, Creelman R. Mutualism between the carnivorous purple pitcher plant and its inhabitants. The American Midland Naturalist. 1984;112(2):294–304.

69. Bernardin J, Young E, Gray S, Bittleston L. Bacterial community function increases leaf growth in a pitcher plant experimental system. mSystems. 2024;9:e01298–24.

70. Luciano C, Newell. Effects of prey, pitcher age, and microbes on acid phosphatase activity in fluid from pitchers of *Sarracenia purpurea* (Sarraceniaceae). PLoS One. 2017;12(7):e0181252. doi:doi:%2010.1371/journal.pone.0181252

71. Miller T, Cassell D, Johnson C, et al. Intraspecific and interspecific competition of *Wyeomyia smithii* (Diptera: Culicidae) in pitcher plant communities. American Midland Naturalist. 1994;131.

72. Wakefield A, Gotelli N, Wittman S, Ellison A. The effect of prey addition on nutrient stoichiometry, nutrient limitation, and morphology of the carnivorous plant *Sarracenia purpurea* (Sarraceniaceae). Ecology. 2005;86:1737–1743.

73. Carmickle R, Horner J. Impact of the specialist herbivore *Exyra semicrocea* on the carnivorous plant *Sarracenia alata*: a field experiment testing the effects of tissue loss and diminished prey capture on plant growth. Plant Ecology. 2019;220(6). doi:DOI:%2010.1007/s11258-019-00935-y

74. Klecka J, Hadrava J, Kolouskova. Vertical stratification of plant-pollinator interactions in a temperate grassland. Peer J. 2018;6:e4998. doi:doi:%2010.7717/peerj.4998.

75. Beikmohamadi L. Effects of Hybridization on Phytotelma communities in *Sarracenia purpurea* and *Sarracenia jonesii*. Capstone: The UNC Asheville Journal of Undergraduate Scholarship. Published online 2018:218–224.

76. Morgan GW. *Sarracenia purpurea* var. *montana* and *Sarracenia jonesii*: A Study of Effects of Hybridization. *Capstone*, The UNC Asheville Journal of Undergraduate Scholarship. 2019;31(2). doi:https://janeway.uncpress.org/capstone/article/id/3320/

77. Irwin D, Schluter D. Hybridization and the Coexistence of Species. The American Naturalist. 2022;200(3):E93–E109. doi:10.1086/720365

78. Lee Y, Braglia L, Stepanenko A, et al. Hybridity of mainly asexually propagating duckweeds in genus *Lemna* – dead end or breakthrough? New Phytologist. 2026;250:629–647. 10.1111/nph.70748

79. Thompson K, Brandvain Y, Coughlan J, et al. The Ecology of Hybrid Incompatibilities. Cold Spring Harbor Perspectives in Biology. 2024;16(9):a041440

80. Wang D, Xu X, Zhang H, et al. Abiotic Niche Divergence of Hybrid Species from Their Progenitors. The American Naturalist. 2022;200(5):634–645. doi:doi:%2010.1086/721372

81. Uzzell L. Reproductive Effort and Output in Two Species of *Sarracenia* (Pitcher Plant) and Their Hybrids. *Capstone*, The UNC Asheville Journal of Undergraduate Scholarship. 2017;30(1). doi:https://janeway.uncpress.org/capstone/article/id/3090/

82. Ellison A, Parker J. Seed dispersal and seedling establishment of Sarracenia purpurea (Sarraceniaceae). 2002;89(6):1024–1026. doi:doi:%2010.3732/ajb.89.6.1024

83. Waser N, Chittka L, Price M, Williams N, Ollerton. Generalization in pollination systems, and why it matters. Ecology. 1996;77:1043–1060. 10.2307/2265575

84. Nomura Y, Shimono Y, Mizuno N, et al. Drastic shift in flowering phenology of F1 hybrids causing rapid reproductive isolation in *Imperata cylindrica* in Japan. Journal of Ecology. 2022;110(7):1548–1560. 10.1111/1365-2745.13890

85. Berry EJ, Cleavitt NL. Population dynamics and comparative demographics in sympatric populations of the round-leaved orchids *Platanthera macrophylla* and *P. orbiculata*. Population Ecology. 2021;63(4):274–289. doi:10.1002/1438-390X.12092

